# Proteasome homeostasis is essential for a robust cauliflower mosaic virus infection

**DOI:** 10.1101/2021.03.24.436740

**Authors:** Aayushi Shukla, Suayib Ustun, Anders Hafrén

**Affiliations:** Department of Plant Biology, Uppsala BioCenter, Swedish University of Agricultural Sciences and Linnean Center for Plant Biology, Box 7080, 75007 Uppsala, Sweden; University of Tübingen, Center for Plant Molecular Biology (ZMBP), Tübingen, Germany

## Abstract

The ubiquitin-proteasome system (UPS) is essential for the maintenance and shifts in protein homeostasis, and thereby forms a founding pillar in virtually all cellular processes including plant immunity and viral infections. According to its importance in fine-tuning the complex plant immune response, proteasomal defects result in divergent outcomes including both resistance and susceptibility phenotypes in response to viruses. The final outcome will largely depend on the specific virus and its specific co-adaptation with the UPS as well as the immune system. Here, we show that cauliflower mosaic virus (CaMV) relies on the proteasome for robust infection. The proteasome system is induced during infection via SA and supports systemic accumulation of the virus as well as plant growth performance during infection. This establishes the UPS as a win-win pathway for the plant and the virus, and together with our demonstration of a proteasome-suppressing viral effector, the intimacy between the proteasome and CaMV is fortified.

## Introduction

The ubiquitin-proteasome system (UPS) is an essential part of protein homeostasis and represents the major protein catabolic pathway alongside autophagy in eukaryotic cells (Vierstra, 2009). In addition to degrading malfunctional proteins, the UPS provides cells with an efficient machinery to adjust numerous cellular responses by rapidly degrading regulatory proteins in a highly selective manner. Multiple studies have together revealed that the UPS regulate plant homeostasis in several contexts, including plant development, cell division, endoreduplication, transcription, phytohormonal responses, perception of environmental cues as well as abiotic and biotic stress interactions (Sadanandom et al., 2012; Xu and Xue, 2019). Indeed, it is valid to view the UPS as a central regulatory hub in most, if not all, major homeostatic shifts occurring when cells undergo reprogramming events in developmental and environmental responses.

The canonical pathway of protein degradation by the UPS involves tagging target proteins with ubiquitin through the action of ubiquitin activating enzymes (E1s), ubiquitin conjugating enzymes (E2s) and ubiquitin ligases (E3s) (Vierstra, 2003). The importance, width and specificity of the UPS becomes evident when looking at the enormous diversity of the E3s in plants, with over 1.400 *in silico* identified members (Gagne et al., 2002; Stone et al., 2005; Vierstra, 2009). Because E3s have a major role in defining the specificity of which proteins are tagged with ubiquitin, the fact that over 5% of Arabidopsis genes appear to encode for E3s supports extensive regulative capacity by this node of the UPS. Following ubiquitin tagging, proteins are recognized and degraded by the proteasome within the UPS. The exact composition of proteasomes may show extensive variation (Morozov and Karpov, 2019), but the canonical 26S proteasome of the UPS is a large 2.5 MDa complex consisting of two multi-subunit complexes; the 19S regulatory particle involved in ubiquitin recognition and the 20S catalytic chamber involved in proteolysis. Adding proteasome compositional variation together with E3 ligase specificity, ubiquitin chain diversity and a growing number of ubiquitin-like modifiers (Vierstra, 2012), the full regulative plasticity of the UPS in cellular homeostasis appears ubiquitous.

A common factor for successful plant adaptation to various environmental stresses is a precise cellular reprogramming. In accordance with the global importance of the proteasome in cellular homeostasis, plants with defects in various proteasome components are frequently compromised in stress adaptation and tolerance (Smalle et al., 2003; Kurepa et al., 2008; Xu and Xue, 2019). This includes increased sensitivity to the abiotic stress factors drought, heat and salinity as well as different pathogens. Moreover, even the lack of individual E3s frequently reduce abiotic and biotic stress tolerance, together manifesting that rather the whole UPS and not only the proteasome, is essential for plant reprogramming in response to the environment (Xu and Xue, 2019).

It is established that plant immunity depends on UPS functions in a broader context (Goritschnig et al., 2007; Marino et al., 2012; Langin et al., 2020), likely involving a multitude of regulated plant as well as pathogen components. This dependence is partially reflected by the reduced pathogen resistance in 19S proteasome mutants *rpt2a*, *rpn1a*, *rpn8a*, *rpn12a* and *rpn10* (Yao et al., 2012; Ustun et al., 2016; Ustun et al., 2018). We can expect both shared and specific phenotypes in different proteasome mutants, where dysregulated salicylic acid (SA) responses would be a general cause of altered pathogen resistance. SA is a master regulator of plant immunity and accumulates both in local and distal systemic tissue upon plant infection with various pathogens (Gao et al., 2015). Interestingly, *rpt2a*, *rpn1a*, *rpn8a* and *rpn12a* all show reduced SA accumulation and/or expression of the SA-defense hallmark gene *PR1* upon infection with pathogenic bacteria (Yao et al., 2012; Ustun et al., 2016). *PR1* expression is largely mediated through the SA master-regulator NPR1 transcription factor, in a process proposed to involve a continuous turn-over of NPR1 through the proteasome (Spoel et al., 2009). In addition to local defense, SA accumulates and promotes systemic acquired resistance in distal tissues (SAR) through NPR1 in response to various pathogens (Dong, 2004; Fu and Dong, 2013; Gao et al., 2015). The other way around, SA has also been identified as an inducer of the proteasome (Gu et al., 2010; Ustun et al., 2013).

There are several discoveries of interplay between plant viruses and the UPS (Alcaide-Loridan and Jupin, 2012; Verchot, 2016). As one example, the genus potyviruses include many findings; strong accumulation of ubiquitinated proteins during infection (Hafren et al., 2018), viral protein HCpro interacting with several different proteasome subunits (Jin et al., 2007) and also proposed to inhibit an antiviral RNAse activity of the proteasome subunit PAE1 (Dielen et al., 2011). Furthermore, the potyviral coat protein is targeted by the E3 ligase CHIP for proteasomal degradation to promote virus replication (Hafren et al., 2010; Lohmus et al., 2017). We expect that this kind of multiplexity of interactions observed for potyviruses and the UPS, is more likely common than exceptional for most plant viruses and thereby that there is a yet largely uncovered plethora of UPS mechanisms operating in plant virus interactions.

In the current study, we have analysed the role of the proteasome system in cauliflower mosaic virus (CaMV) infection. We found that the proteasome activity is induced by SA during infection, but independent of NPR1 as previously observed for bacteria. We further demonstrate that viral effector protein P6 can suppress proteasome activity, suggesting close interplay between the proteasome and the virus. Finally, we show that systemic accumulation of CaMV is hampered in plants with defects in the proteasome, together establishing that a properly tuned proteasome system is elementary for a robust CaMV infection.

## Results

### Proteasome activity is induced during CaMV infection

Our first goal was to determine whether the proteasome and ubiquitin-system responds to CaMV infection. For this we infected Arabidopsis WT plants with CaMV and estimated proteasome activity based on fluorescence release from the substrate Succ-LLVY-AMC in lysates prepared 21 days after infection (DAI) (Figure 1A). This activity was significantly increased in CaMV infected tissue compared to the uninfected control, suggesting enhanced proteasome activity. Several transcripts encoding for proteasome subunits are part of the proteasome regulon that responds e.g. to the proteasome inhibitor MG132 (Gladman et al., 2016), including the highly responsive *Rpt2a* while *Rpt2b* is less responsive. RT-qPCR analysis showed that the transcript level of *Rpt2a* was elevated during CaMV infection while the impact on *Rpt2b* was comparably marginal (Figure 1B). Overall, this response phenocopied the proteasome inhibitor MG132 albeit the magnitude of the response was less (Figure 1C), supporting proteasome activation during CaMV infection. To estimate the overall functionality of the UPS during CaMV infection, we also compared the level of high-molecular weight ubiquitin present in infected and control tissues using ubiquitin western blotting analysis (Figure 1D). The ubiquitin signal appeared stronger in CaMV lysates, but this difference was minor compared to that observed in response to the proteasome inhibitor MG132 (Book et al., 2010) or *Turnip mosaic virus* (Hafren et al., 2018). Nevertheless, a GFP-ubiquitin (Ub) expressing marker line also supported some elevation in ubiquitin levels and revealed the formation of some ubiquitin-positive cytoplasmic foci in the presence of CaMV (Figure 1E). In mock, GFP-Ub was faintly detectable. Taken together, we conclude that CaMV infection induces proteasome activity, with a concurrent modest accumulation of high-molecular weight ubiquitin.

**Figure 1.**
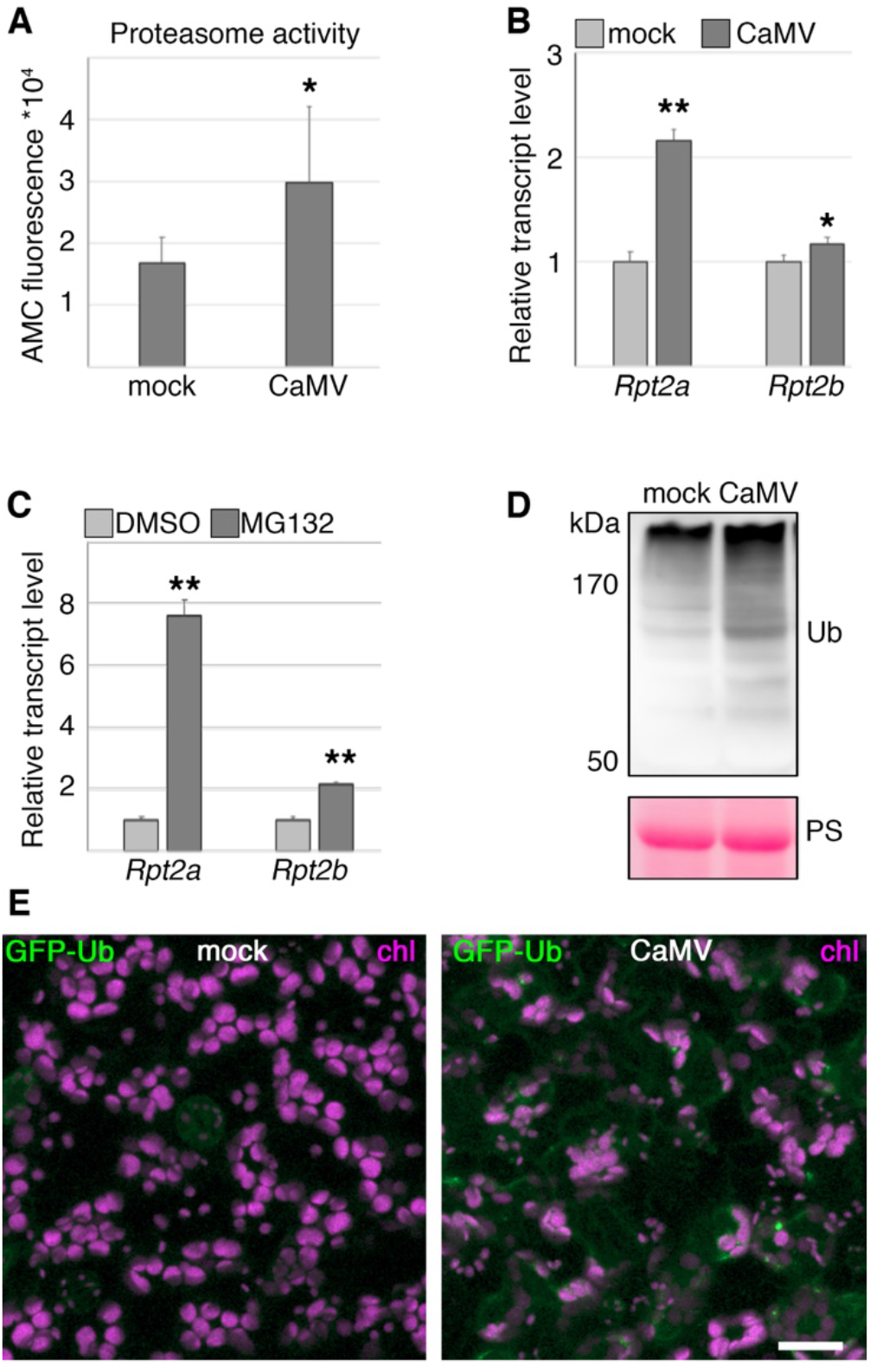
CaMV infection activates the proteasome and ubiquitin responses A) Proteasome activity measured from mock and CaMV infected plants at 21 DAI. The proteasome activity was measured using the substrate Succ-LLVY-AMC according to absolute fluorescence and its kinetics. *n*=4. B and C) Transcript levels for *RPT2a* and *RPT2b* in mock and CaMV infected plants at 21 DAI (B) or DMSO and MG132 treated seedlings (C) were determined by RT-qPCR. *PP2a* was used as reference. *n*=4. D) Western blot detection of ubiquitin (Ub) from mock and CaMV infected plants at 21 DAI. Ponceau S (PS) staining shows the loading. E) Confocal images of an Arabidopsis marker-line stably expressing GFP-tagged ubiquitin (GFP-Ub in green) in control and during CaMV infection at 21 DAI. Chloroplasts (chl) are shown in magenta. Scale bar = 20 μm. Statistical significance by Students *t*-test; * *p*<0.05, ** *p*<0.01.

### Viral protein P6 suppresses proteasome activity induced by salicylic acid during CaMV infection

The increased proteasome activity observed during CaMV infection (Figure 1A), could be part of an infection response regulated through some of the major antiviral defense pathways in plants. Therefore, we analyzed whether proteasome activation depended on either salicylic acid (SA) or RNA silencing pathways by infecting plants expressing SA-degrading bacterial *NahG*, SA-signaling mutant *npr1* as well as RNA silencing triple mutants *dcl2 dcl3 dcl4* and *rdr1 rdr2 rdr6*. Proteasome activity measurements revealed that the increase observed in WT was identical only in *npr1*, while being highly compromised in plants expressing *NahG* (Figure 2A). The RNA silencing mutants showed a significant increase in proteasome activity in contrast to *NahG*, but the fold of increase was lower than in WT and *npr1* especially for *rdr1 rdr2 rdr6*. Because viral DNA levels were comparable between the different genetic backgrounds, the absence and reduction of proteasome activation in *NahG* and *rdr1 rdr2 rdr6*, respectively, was unlikely linked to altered infection levels (Figure 2B). Furthermore, the SA synthesis mutant *sid2* did not either show elevated proteasome activity in response to CaMV (Figure 2C). This established that SA was essential for increasing proteasome activity during CaMV infection. However, the NAC53/NAC78 transcription factors that drive the proteasome transcriptional regulon in response to the inhibitor MG132 (Gladman et al., 2016), were not essential for infection to induce proteasome activity (Figure 2C), suggesting that induction by CaMV goes through some other pathway. Likewise, we found that *Pseudomonas syringae DhrcC*, which is not able to deliver type-III effector proteins and previously shown to induce proteasome activity in an NPR1-dependent manner, did not depend on the NAC transcription factors to induce proteasome function (Figure 2D).

**Figure 2.**
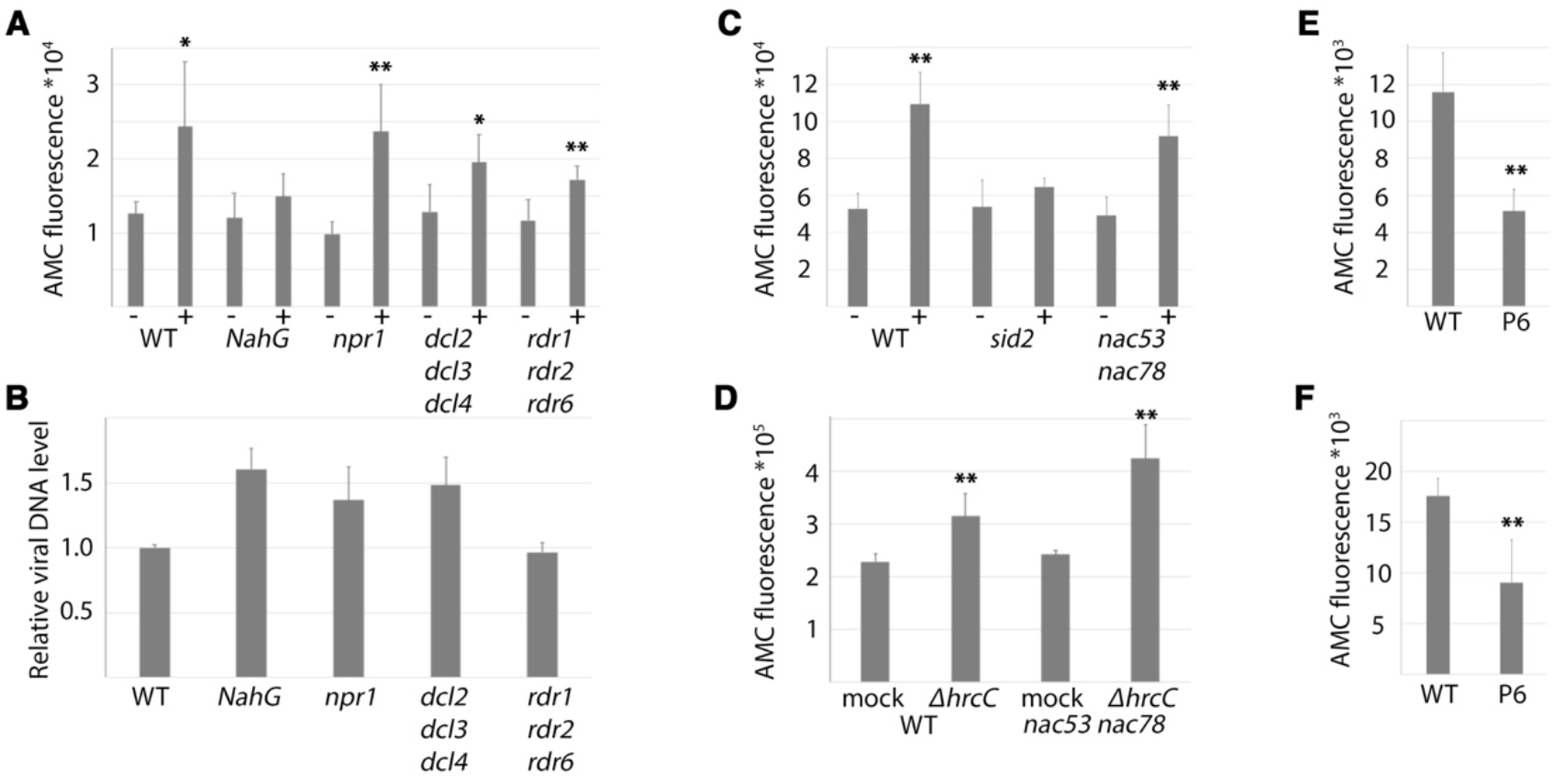
CaMV-induced proteasome activity requires salicylic acid A) Proteasome activity was measured using the Succ-LLVY-AMC fluorescence release assay in mock (-) and infected (+) WT, *NahG*, *npr1*, *dcl2 dcl3 dcl4* and *rdr1 rdr2 rdr6* plants at 21 DAI. *n*=4. B) Viral DNA levels were determined by qPCR in parallel with proteasome activity measurements in (A). rDNA was used as reference. *n*=4. C) Proteasome activity as (A) in mock and infected SA-biosynthesis mutant *sid2* and proteasome regulon mutant *nac53 nac78*. *n*=4. D) Proteasome activity in mock and *ΔhrcC* infected proteasome regulon mutant *nac53 nac78*. *n*=4. E) Proteasome activity in Arabidopsis WT and P6-mRFP transgenic 4-week-old plants. *n*=4. F) Proteasome activity in *Nicotiana benthamiana* plants 3 days after infiltration of mRFP or P6-mRFP. *n*=4. Statistical significance by Students *t*-test, * *p*<0.05, ** *p*<0.01.

Interestingly, the viral protein P6 was demonstrated to dampen SA-responses in a mechanism involving NPR1 (Love et al., 2012). Additionally, P6 was reported to inhibit SA-dependent activation of autophagy (Zvereva et al., 2016). Because proteasome activity can be induced by SA directly (Gu et al., 2010; Ustun et al., 2013) and occurs in an SA-dependent manner during CaMV infection, we reasoned that viral P6 is a potential suppressor of the proteasome. Indeed, the proteasome activity was significantly lower in non-infected transgenic Arabidopsis plants expressing a P6-mRFP fusion (Figure 2E). Likewise, transient expression of P6-mRFP in leaves of *Nicotiana benthamiana* caused a significant reduction in proteasome activity levels compared to the mRFP control (Figure 2F). These results show that CaMV infection induces proteasome activity via SA in an NPR1-independent manner, and that the viral protein P6 is capable of suppressing the proteasome, together suggesting that the proteasome may be a central component of CaMV disease that both the plant and the virus are regulating.

### Proteasome functionality influences CaMV disease development

The above results suggested that the proteasome system responds to infection, and next we set out to determine whether the proteasome influences the severity of CaMV disease. A proteasome knock-out is lethal, but several viable T-DNA knock-out mutants for individual proteasome subunits have been described that show both common and specific plant phenotypes as well as proteasome defects. The *pbe1* (20S β5 subunit) mutant was recently shown to have proteasome assembly and ubiquitin degradation defects upon salt stress, and the mutant is sensitive to both DTT and MG132 (Han et al., 2019). The *pae1* (20S α5 subunit) mutant shows enhanced susceptibility to a potyvirus, potentially linked to PAE1 RNAse activity (Dielen et al., 2011), but whether proteasomes are compromised in *pae1* plants is not known. Notably, *pae1* is unlikely a strong proteasome mutant as it lacks the evident morphological alterations seen for *pbe1*, *rpn12a* and *rpn10*. The *nac53 nac78* transcription factor double mutant fails to up-regulate the transcriptional regulon of proteasome subunits and shows sensitivity to MG132 (Gladman et al., 2016), but does not regulate CaMV-induced proteasome activity (Figure 2) and is not expected to have directly compromised proteasome functions. Both *rpn12a* and *rpn10* 19S mutant proteasomes show decreased 26S-dependent (ubiquitinated proteins) but increased 20S-dependent (oxidized proteins) degradation and various developmental alterations (Smalle et al., 2002; Smalle et al., 2003; Kurepa et al., 2008). Furthermore, *rpn10* is so far the only demonstrated proteasome mutant that clearly accumulates ubiquitinated proteins under normal conditions (Smalle et al., 2003), likely in part owing to its functions as the receptor for ubiquitinated proteasome degradation by autophagy (Marshall et al., 2015). Together, this set of mutants should cover a range of proteasome dysfunctions.

When this collection of proteasome mutants was infected with CaMV, we found that all mutants except for *nac53 nac78* showed significantly increased biomass loss compared to WT plants by 21 DAI (Figure 3A). Interestingly, the biomass loss phenotype recovered by 35 DAI in *pbe1* and *pae1*, but remained significantly reduced in *rpn12a* and *rpn10* (Figure 3B). The uninfected control plants did not show any significant differences in biomass at either time-points (Figure S1). A representative image of control and infected plants at 35 DAI shows additionally that older leaves of *rpn12* and especially *rpn10* senesced pre-maturely during CaMV infection (Figure 3C). Because SA can slow growth and promote leaf senescence, we tested whether SA responses were altered in these mutants. Indeed, expression of the SA-responsive marker gene *PR1* was highly increased in *rpn12a* and *rpn10* mutants compared to the other mutants and WT (Figure 3D). This suggests that *rpn12a* and *rpn10* have elevated SA responses in response to CaMV, which could explain their enhanced senescence and growth reduction. Together, these results show that proteasome functions counteract plant biomass loss and SA responses in CaMV infection.

**Figure 3.**
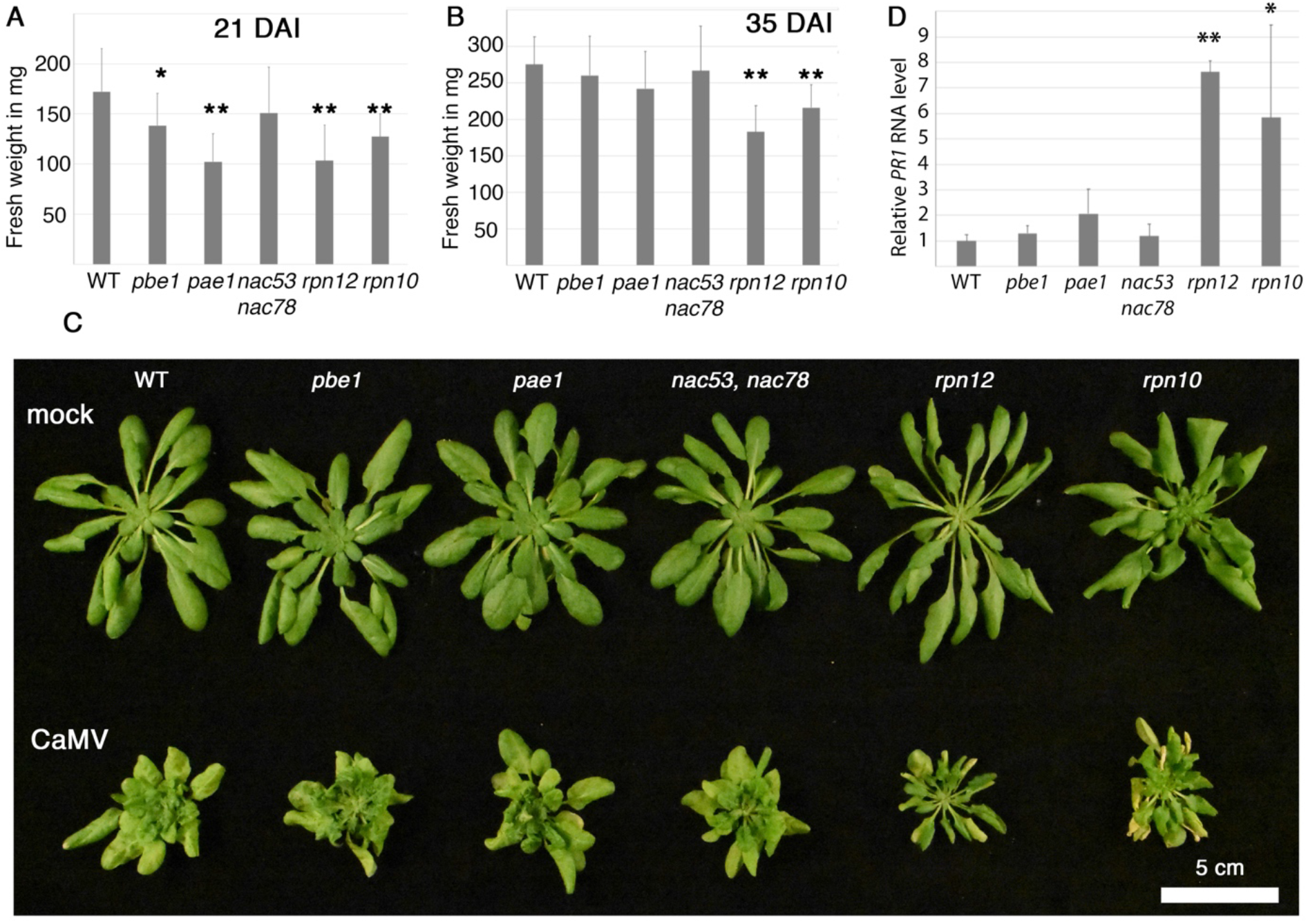
Defects in the proteasome reduce plant vigor in CaMV disease A and B) The fresh weight of CaMV infected WT, *pbe1, pae1, nac53 nac78, rpn12a* and *rpn10* plants at 21 DAI (A) and 35 DAI (B). *n*>8. C) Image of plants representative of infection (lower row) and uninfected control plants (upper row) at 35 DAI. D) *PR1* expression determined by RT-qPCR in CaMV infected plants at 21 DAI. *PP2a* was used as reference. *n*=4. Statistical significance by Students *t*-test; * *p*<0.05, ** *p*<0.01.

### The proteasome supports systemic CaMV accumulation

Very interestingly, the youngest leaves of especially *rpn10* and *rpn12a* lacked the typical vein-clearing and rosette morphology distortion that was evident in infected WT, *pae1* and *nac53 nac78* plants at 35 DAI (Figure 3C). The symptoms appeared somewhat attenuated in the *pbe1* mutant as well, but not as clearly as in *rpn10* and *rpn12a*. As the first quantitative measure of this phenotype, we calculated the ratio of asymptomatic to symptomatic leaves per plant as defined by presence of vein-clearing to support our initial observation (Figure 4A). A detailed picture clearly shows the absence of this symptom in the upper leaves of *rpn10* compared with WT (Figure 4B).

**Figure 4.**
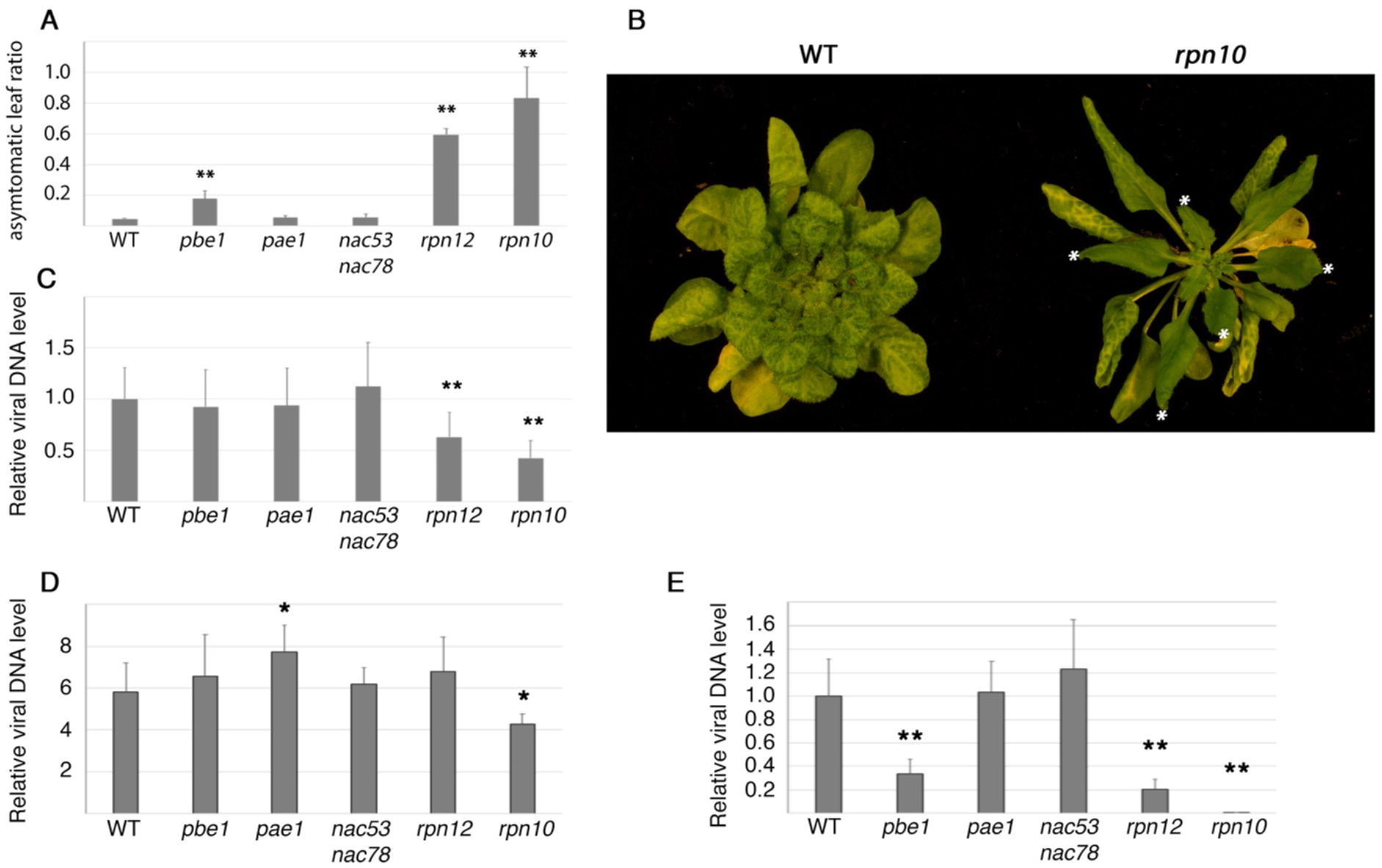
The proteasome supports systemic accumulation of CaMV A) The ratio between leaves without and with vein-clearing symptoms per plant at 35 DAI. *n*=5. B) A representative picture of infected WT and *rpn10* plants at 40 DAI, where the largely asymptomatic younger leaves in the rosette are marked with a white asterisk. C)Viral DNA levels determined by qPCR at 21 DAI. rDNA was used as reference. *n*=8. D and E) Viral DNA levels were determined separately in the lower (D) and uppermost (E) part of the plant rosette at 35 DAI. *n*=4. Statistical significance by Students *t*-test; * *p*<0.05, ** *p*<0.01.

Our next interest was to see whether virus accumulation followed the symptom severity. We inoculated WT, *pbe1*, *pae1*, *nac53 nac78*, *rpn12a* and *rpn10* plants and analyzed virus accumulation at 21 DAI (Figure 4C), revealing significantly reduced levels of viral DNA in both *rpn12a* and *rpn10* plants. Notably, the decrease in virus accumulation was consistently stronger in *rpn10* than *rpn12a*. At 35 DAI, when the symptom attenuation phenotype was more evident (Figure 4B), we analyzed the level of viral DNA separately from the lower part of the plant rosette and the upper-most six leaves. In the lower part of the plant, all mutants showed the typical vein-clearing symptom of CaMV and viral DNA accumulated to comparable levels even though still somewhat reduced in *rpn10* (Figure 4D). However, viral DNA was strongly reduced in the upper leaves of *pbe1* and *rpn12a*, and close to absent in *rpn10* (Figure 4E). These results strongly support that systemic virus accumulation is compromised in *pbe1*, *rpn12a* and *rpn10* mutants. Furthermore, the robust accumulation of viral DNA in the lower leaves support that CaMV is fully capable of strong replication in these mutants, pointing towards enhanced systemic resistance rather than a direct effect on virus replication. As the *rpn10* mutant is a more severe proteasome mutant than *rpn12a* (Kurepa et al., 2008), we suspect that the reduction on CaMV accumulation may correlate with the severity of the mutation where *rpn10* is worst, *pbe1* and *rpn12a* are in the middle and *pae1* and *nac53 nac78* are close to WT.

## Discussion

A functional proteasome is important for proper plant responses and tolerance to several abiotic, biotic and biochemical stress conditions, including salinity, drought, heat, UV, micronutrient imbalance (Zn, Br), oxidative stress, proteasome inhibition, ER stress and bacterial disease (Xu and Xue, 2019). The proteasome regulates many plant defense components in conjunction with the ubiquitin system, including SA-induced NPR1 that promotes SAR and *Pathogenesis-Related* (*PR*)-gene expression (Spoel et al., 2009). Similar to bacterial diseases (Hatsugai et al., 2009; Yao et al., 2012; Ustun et al., 2016), the proteasome is likely important for mounting a robust defense response during virus infections by direct regulation of plant immunity. Viruses as intracellular pathogens are furthermore directly exposed to potential regulation by the proteasome and likewise, the proteasome is readily available for exploitation and manipulation by viruses. Indeed, numerous studies have together outlined an intimate and many times highly individual interaction between the UPS and diverse animal viruses (Luo, 2016; Tang et al., 2018). Their proviral interactions include degradation of antiviral host proteins, dampening the immune response, fine-tuning viral protein levels and viral protein activation through ubiquitin-like modifiers that collectively cover all major steps of infection viral cycles. Fundamentally corresponding interactions have also been identified for plant viruses (Dielen et al., 2010; Alcaide-Loridan and Jupin, 2012; Verchot, 2016). The plant UPS and the proteasome is known to destroy viral proteins (Reichel and Beachy, 2000; Barajas et al., 2009), optimize viral protein levels (Drugeon and Jupin, 2002; Hafren et al., 2010; Chenon et al., 2012; Lohmus et al., 2017; Fieulaine et al., 2020), degrade the antiviral RNA silencing component AGO1 (Chiu et al., 2010), be a point of suppressing immunity (Liu et al., 2002; Eini et al., 2009; Lozano-Duran et al., 2011; Jia et al., 2016) as well as the proteasome being directly targeted by a viral protein (Ballut et al., 2005; Jin et al., 2007; Dielen et al., 2011). Thus, the UPS establishes a ubiquitous pro- and antiviral system in both plant and animal virus diseases. Together with the more recently emerged divergent roles of autophagy (Kushwaha et al., 2019), protein catabolism in general stands a central position in plant virus disease.

Our present work identifies several aspects of proteasome importance in CaMV disease; i) the proteasome activity is induced during infection via SA, ii), the viral protein P6 can suppress proteasome activity, iii), plant growth during infection is compromised in proteasome mutants and, iv), several proteasome mutants show reduced systemic infection. Currently, surprisingly little is known about how the level of proteasome activity is regulated in plants. At the transcriptional level, NAC53 and NAC78 transcription factors drive the proteasome stress regulon at least in response to the proteasome inhibitor MG132 (Gladman et al., 2016), but they were not important for proteasome up-regulation during either CaMV or *Pseudomonas syringae ΔhrcC* infections. Instead, we identify SA as an essential component driving proteasome induction during CaMV infection. Indeed, SA alone can increase proteasome activity upon foliar application (Ustun et al., 2013) and both *Pseudomomas syringeae* and *Xanthomonas campestris* modulates proteasome activity through the SA-regulator NPR1 (Ustun et al., 2013; Ustun et al., 2016), while in the case of CaMV this is fully NPR1 independent. We also found that expression of the SA-marker gene *PR1* is strongly up-regulated in the *rpn12a* mutant during CaMV infection, being in sharp contrast to what was observed for *Pseudomomas* (Ustun et al., 2016). Together, these findings suggest that the connective mechanisms between SA and the proteasome are different for extracellular pathogenic bacteria and CaMV. Nevertheless, all of them have suppressive effectors as we now found that also the viral P6 protein can reduce proteasome activity. This is likely connected to the ability of P6 to inhibit SA-dependent signaling (Love et al., 2012), thus making it impossible to say whether the primary target is the proteasome or other SA-dependent viral defense responses (Carr et al., 2019; Murphy et al., 2020). In support of a more complex role for P6 in SA regulation, P6 was demonstrated to also inhibit SA-induced autophagy (Zvereva et al., 2016) and autophagy was shown to regulate CaMV (Hafren et al., 2017). As P6 was proposed to regulate SA via NPR1 (Love et al., 2012), one alternative is that the NPR1 route of proteasome activation is shut down by P6 during CaMV infection while an SA-dependent but NPR1-independent mechanisms still functions. This would agree with the analogous proteasome response between the *npr1* mutant and WT. A deeper understanding of the mechanistic underpinnings of SA-dependent proteasome activation by CaMV infection and repression by P6 represents a future challenge.

We used several T-DNA insertion mutants to evaluate the importance of the proteasome on CaMV accumulation and disease. While individual proteasome subunit mutants could all have compromised assembly of the full proteasome and share responses such as sensitivity to MG132 (Smalle et al., 2002; Smalle et al., 2003; Kurepa et al., 2008; Lee et al., 2011; Gladman et al., 2016; Han et al., 2019), they are not analogous in terms of their effect on plant morphology in general or impact on the proteasome. Accordingly, these mutants also respond differently to CaMV. The *nac53 nac78* double-mutant phenocopied WT, conceivable as the NAC’s are uncoupled from CaMV-induced proteasome activity. Plant growth was reduced in all other mutants during CaMV infection. This was not associated with resistance defects, and suggests that a properly tuned proteasome is generally important for plant growth during infection. Furthermore, older leaves of *rpn12a* and *rpn10* showed premature senescence, likely associated with hyperactivation of SA and *PR1* induction in these mutants. Possibly, SA contributes to enhanced systemic resistance against CaMV in proteasome mutants, but this is not supported by *pbe1* that does not show similar *PR1* activation as *rpn12a* and *rpn10*. The proteasome system is frequently co-opted for fine-tuning viral protein levels (Luo, 2016; Tang et al., 2018), and the capsid protein of CaMV was identified to be a substrate for the proteasome (Karsies et al., 2001) and also autophagy (Hafren et al., 2017). Misregulated capsid protein levels are potential hazards for viruses due to their capacity to interfere with various steps of the infection cycle (Ivanov and Makinen, 2012). However, because CaMV accumulates normally in older systemic leaves irrespective of proteasome defects, we propose that the reduction of virus in young systemic leaves is indirect immunity and not related to capsid protein degradation. Our current working model based on the presented results is that the systemic accumulation of CaMV depends on a functional proteasome, and the degree of proteasome defect in the mutants correlate with CaMV accumulation. The strong connectivity between UPS and plant immunity in general could suggest that a properly regulated immune system by the proteasome is elementary for CaMV (Goritschnig et al., 2007; Marino et al., 2012; Ustun et al., 2016). We conclude that the two major protein catabolic pathways, proteasome and autophagy, are both tightly nested in CaMV infection but with highly different roles (Hafren et al., 2017), and leave a future challenge to determine which defense pathway is activated to provide resistance against CaMV in proteasome mutants – RNA silencing, SA, autophagy or other pathways. A hypothetical model connecting the proteasome, SA and CaMV infection is presented in (Figure 5).

**Figure 5.**
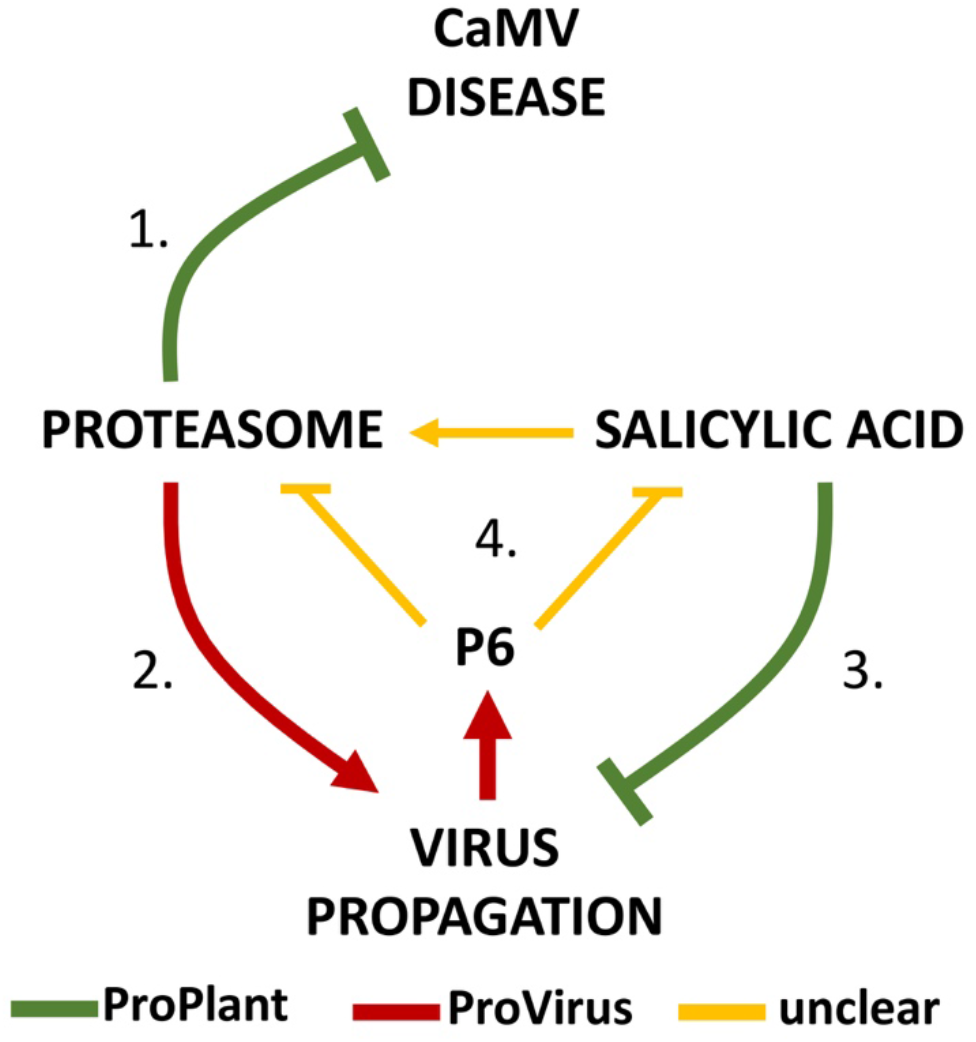
A model for the interconnections between proteasome, SA and CaMV infection Disruption of proteasome homeostasis results in increased disease detected as increased plant biomass loss and senescence (1), possibly involving escalated SA-responses. Notably, this is uncoupled from resistance as the proteasome also supports virus accumulation (2), outlining the proteasome as ProVirus-ProPlant. In contrast, SA provides resistance against CaMV by restricting virus accumulation (3) but also drives proteasome up-regulation (4). Together with the capacity of viral P6 to suppress both SA and proteasome activity, the connectivity between CaMV, the proteasome and SA is unlikely linear but rather a complex network of interactions that fine-tune several distinct infection processes.

## Methods

### Plant material, growth conditions and reagents

Arabidopsis T-DNA insertion mutants were *pbe1* (SALK_ 053781) (Huang et al., 2006), *pae1* (SALK_ 151939) (Dielen et al., 2011), *nac53-1 nac78-1* (SALK_009578C; SALK_025098) (Gladman et al., 2016), *rpn12a-1* (Smalle et al., 2002) and *rpn10-1* (Smalle et al., 2003). The slower germination of *rpn10-1* was adjusted for by sowing these seeds 3 days in advance. The Arabidopsis line expressing GFP-ubiquitin was described in (Hafren et al., 2018). The P6 ENTRY clone was described (Hafren et al., 2017) and recombined into pGWB654 (Nakagawa et al., 2007) for transient expression and transformation of Arabidopsis. Arabidopsis plants were grown on soil for infection experiments under short-day conditions (8/16-h light/dark cycles) in a growth cabinet, and *Nicotiana benthamiana* plants were cultivated for transient expression assays under long-day conditions (16/8-h light/dark cycles) in a growth room at 150 μE/m^2^s, 21°C, and 70% relative humidity, respectively.

### Western blot analysis

For ubiquitin detection, plant tissue was ground in protein extraction buffer (100 mM Tris pH 7.5 and 2% SDS) and supplemented with Laemmli sample buffer before boiling 5 min. The cleared extracts were run on an 8% SDS-PAGE gel, transferred to PVDF membrane, blocked with 2% BSA in PBS-T and incubated with 1:1000 diluted polyclonal anti-Ub from Agrisera (AS08 307) followed by secondary anti-rabbit incubation and ECL detection.

### CaMV infection, quantitation and RT-qPCR

The first true leaves of 3-week-old Arabidopsis plants were agroinoculated with CaMV strain CM1841 as before (Hafren et al., 2017). For DNA extraction, plants were sampled in biological replicates, each containing 3 individual plants from which inoculated leaves were removed. DNA was isolated by extracting 50 mg leaf material in 400 *μ*1 250 mM Tris pH 7.5, 5 mM EDTA, 20 mM KC1, 2% SDS. The lysate was heated at 65°C for 30 min, supplemented with 0,2 mg protease K and incubated o/n +37°C, inactivated at 65°C for 30 min and cleared by centrifugation at 17 000 X *g* for 10 min. DNA was precipitated and pelleted from the supernatant with isopropanol 1:1 and 17 000 X *g* for 20 min, followed by washing with 70% EtOH and dissolved in water for qPCR. Quantitative PCR analysis (qPCR) was performed using primers for CaMV and reference ribosomal DNA as described (Hafren et al., 2017). For RNA transcript analysis, RNA was isolated using the Qiagen plant easy kit with on-column DNAse I treatment. cDNA was synthesized with Maxima First Strand cDNA synthesis kit and transcript levels determined by qPCR using primers for *RPT2a* (F_GGAGCGTCGTATGAAGGTGACA, R_GGAGGAGATCAGAAAACATGATC), *Rpt2b* (F_AGAGCGTCGGATGAAAGTGAGC, R_GGCAAACGATTTTATATGATCAG) and *PP2a* as reference (F_ TAACGTGGCCAAAATGATGC, R_GTTCTCCACAACCGCTTGGT).

### Proteasome activity measurement

Proteasome activity was measured essentially as described in (Ustun et al., 2016). Briefly, 3 volumes of buffer (100 mM KHPO_4_ pH 7.5, 5% glycerol, 4 mM ATP, 2 mM DTT) per weight of plant tissue was used to prepare lysates by grinding followed by centrifugation 17 000 X *g* +4°C. 20 *μ*1 of the soluble proteasome extract was used in 100 *μ*1 reactions containing 50 *μ*M Succ-LLVY-AMC proteasome substrate in 50 mM KHPO_4_ pH 7.5 buffer. The kinetics of fluorescence release was recorded from 0 to 120 min, from which a linear range was subsequently selected and used to calculate the total proteasome activity as product between the average fluorescence and the kinetic slope.

### Bacterial infection assay

*Pseudomonas syringae* pv *tomato* DC3000 *ΔhrcC* was grown in King’s B medium with appropriate antibiotics at 28°C. For proteasome activity assays plants were infiltrated with an OD_600_ of 0.1 and proteasome activity was measured 24 hpi.

## Acknowledgement

We acknowledge funding from FORMAS (grant number 2016-01044) for A.H. SÜ was supported by an Emmy Noether Fellowship GZ: UE188/2-1 from the Deutsche Forschungsgemeinschaft (DFG) and through the collaborative research council 1101 (SFB1101). We thank Jan Smalle for kindly sharing *rpn10* and *rnp12a* mutants and Richard D. Vierstra for the *nac53-1 nac78-1* mutant.

---

**Figure S1.**
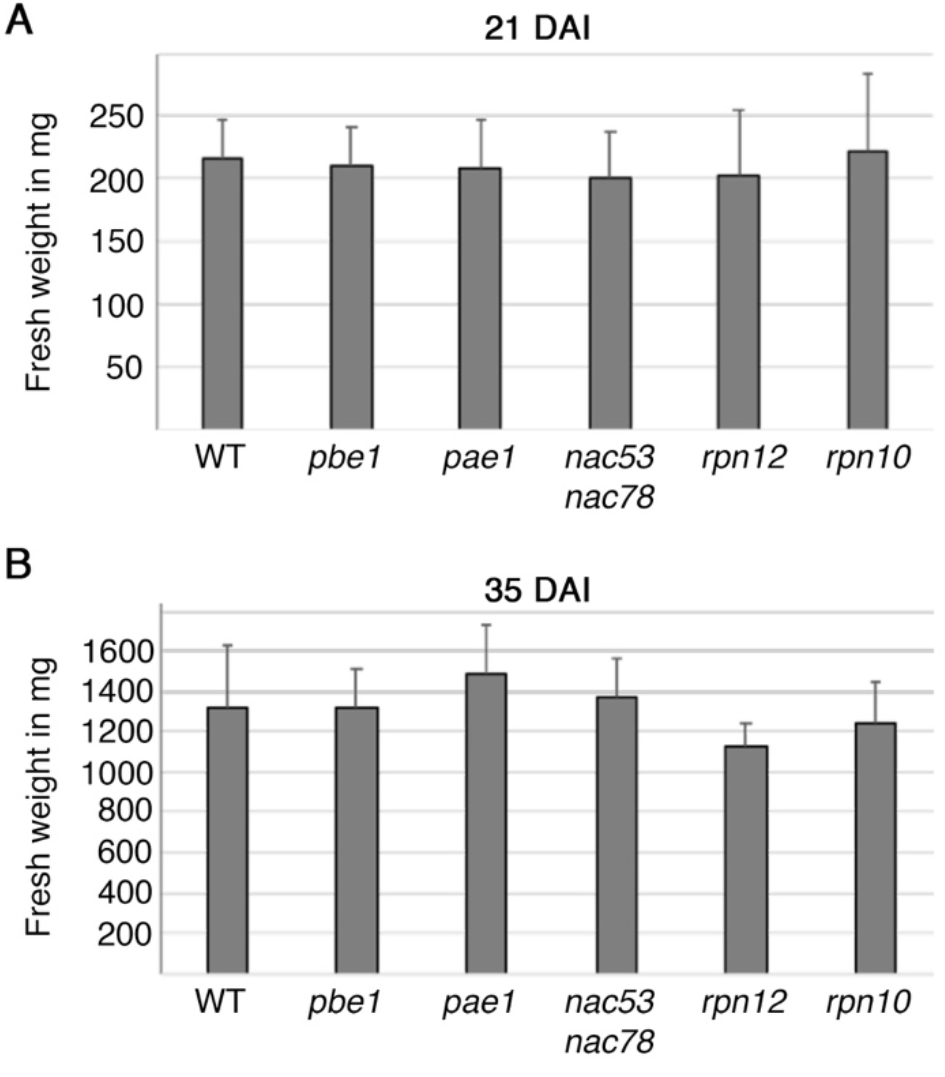
Uninfected proteasome mutants grow WT-like A and B) The fresh weight of uninfected control plants do not usually show significant differences for the proteasome mutants at 21 DAI (A) and 35 DAI (B) in our short-day growth condition. Note that the *rpn10* mutant is put out 3 days earlier than others.

